# Harvesting more reads from single-cell combinatorial barcoding data with *scarecrow*

**DOI:** 10.1101/2025.10.17.683067

**Authors:** D. Wragg, E. Kang, M.D. Morgan

## Abstract

**Summary:** Combinatorial barcoding technologies for single-cell nucleotide sequencing, such as split-pool ligation protocols, involve sequential rounds of cell barcoding to uniquely tag individual cells. The rapid adoption of combinatorial barcoding in recent years is due in part to its scalability across cells and samples. However, small shifts in barcode positions within sequencing reads caused by technical artifacts, e.g. during barcode incorporation or synthesis, can impact the accurate assignment of reads to cell barcodes. Existing processing tools typically assume barcodes contain fixed-length nucleotide sequences located at fixed positions within reads, overlooking any positional variability. Consequently, reads containing truncated or mispositioned barcodes are discarded during initial data processing steps leading to significant data loss. To solve this limitation and maximise the retention of sequencing reads from single-cell combinatorial barcoding experiments, we introduce *scarecrow*. Our tool screens a subsample of reads to generate position-specific barcode profiles, which are then used to flexibly identify barcode sequences in each read whilst accounting for positional errors, a phenomenon we refer to as ‘jitter’. Barcode matches are then prioritised to minimise nucleotide mismatches and the degree of jitter. These initial profiles are subsequently used to extract and error correct barcode combinations in high throughput sequencing libraries. By incorporating jitter into barcode error correction, *scarecrow* enables greater data recovery and improved downstream single-cell analyses. *Scarecrow* is fully open access, implemented in Python, and generates output files using standardised sequence file formats for maximal interoperability. A detailed explanation of the *scarecrow* workflow can be found in the supplementary materials.

## 1 Introduction

Single-cell RNA sequencing (scRNAseq) using combinatorial barcoding strategies such as split-pool ligation offer exponential increases in cell indexing capacity compared to single-indexing strategies (Rosenberg *et al.*, 2018). For both single- and combinatorial-indexing approaches, faithful transcriptome profiling relies on accurate demultiplexing of sequencing reads to single-cell barcodes. One challenge of this demultiplexing step is variability in barcode positioning (e.g. 1-2 bp shifts) or ‘jitter’ due to technical artifacts. This can result from incomplete barcode synthesis (Sun *et al.*, 2025), or be introduced during library preparation and sequencing for example through polymerase slippage during PCR amplification steps, adapter misalignment, or rare indels from enzymatic steps during adapter ligation or tagmentation. Such positional jitter may be more consequential for combinatorial barcoding than single-indexing strategies because of the reliance on multiple barcoding reactions. Existing demultiplexing approaches typically work on the basis that barcodes start either at a fixed position or a position relative to a sequence motif such as a linker (Kuijpers *et al.*, 2024). Jitter or truncation will result in the barcode sequence at the expected location not matching a whitelist of known barcodes even if allowing for a small number of nucleotide substitutions. Consequently, these reads are lost to downstream analysis. Furthermore, not accounting for positional jitter of barcodes may result in incorrect barcode assignment compromising transcriptome profile fidelity. For example, a barcode may be identified at the expected position within a sequencing read with a single mismatch, while a perfect match barcode may start one nucleotide up- or downstream of that position. Efforts to introduce a standardised specification for describing sequencing data, detailing the assay, sequence and library structure, such as *seqspec* (Booeshaghi *et al.*, 2024) are commendable, however the tools that leverage such specifications are still limited by fixed barcode position assumptions.

To address the above challenges, we introduce *scarecrow*. First, *scarecrow* screens a subset of reads against barcode whitelists to generate position-specific barcode distributions. These barcode profiles are then used to flexibly match barcodes to an input whitelist whilst accounting for positional jitter, prioritising barcode matches with the fewest mismatches and smallest jitter. Our benchmarks on Parse Evercode and Scale Biosciences data demonstrate up to a 65% increase in usable reads when accounting for jitter. *Scarecrow* requires no prior knowledge of barcode positions within the library structure, and its outputs are structured for compatibility with standard downstream tools (e.g., *kallisto*, STAR, umi-tools), offering a robust solution for maximizing data yield across diverse single-cell applications.

## 2 Barcode matching

The standard *scarecrow* workflow consists of three steps: *seed* barcodes, *harvest* barcode profiles, and *reap* the sequences (Figure 1A; Supplementary Material S1). While both the *seed* and *reap* steps employ the barcode matching strategy that underpins the utility of *scarecrow, seed* only returns exact matches, without accounting for mismatches or jitter, returning a list of barcode matches and their positions. The *harvest* step gathers the barcode positions from the various barcode indices and generates a profile of expected barcode positions for use with *seed*. Finally, the *reap* command identifies barcode matches based on the expected positions in the profile, while accounting for jitter and allowing for a specified number of mismatches.

**Figure 1.**
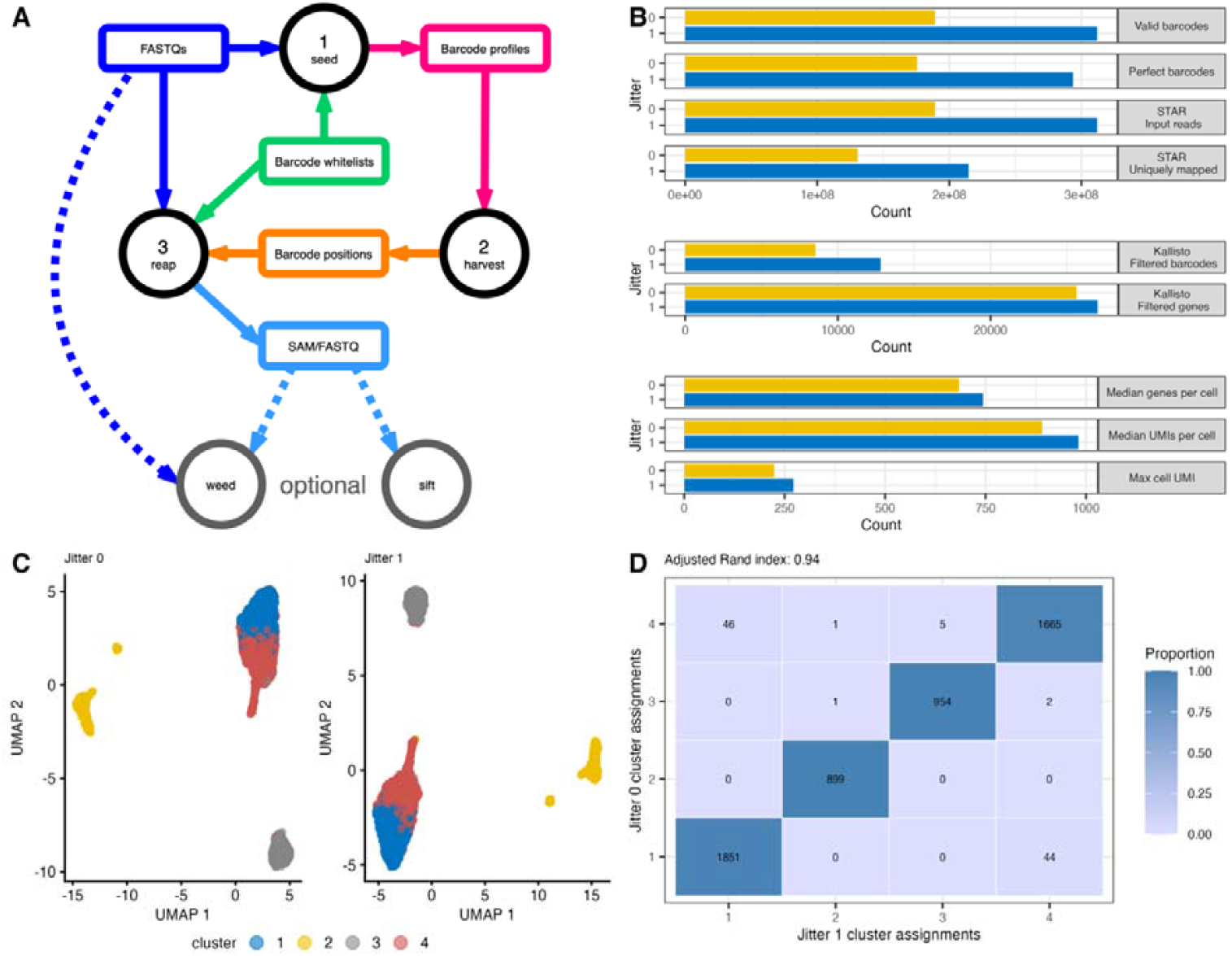
(A) Schematic diagram of the *scarecrow* workflow. This begins with FASTQ files (dark blue box) and barcode whitelists (green box) being passed to *seed*. The resulting barcode profiles (pink box) are passed to *harvest* which outputs a file of expected barcode positions (orange box). These positions, together with the FASTQ files and barcode whitelists are passed to *reap*, which returns either a SAM or interleaved FASTQ file (light blue box) containing the target sequence and barcode information. Optional steps, denoted by dashed lines, include extracting a barcode from the FASTQ read header with *weed*, if the reads have already been demultiplexed on one barcode, and removing any reads containing unmatched barcodes with *sift*. (B) Summary counts from processing Scale Bio data with no jitter and with jitter = 1. This illustrates under each condition: the number of raw reads with valid barcodes, the number of reads with perfectly matching barcodes, the number of reads post-*sift* passed to STAR for alignment, the number of uniquely mapped reads, the number of genes and barcodes remaining post-filtering on a *kallisto* count matrix, the median genes and UMIs per cell in the filtered count matrix, and the max UMI observed across the matrix. (C) Uniform manifold approximation and projections generated from processed and quality controlled ScaleBio profiling of PBMC with no jitter (left) or jitter = 1 (right). Points are coloured by cluster assignment. (D) Heat map indicating agreement of cell cluster assignment from (C) between jitter values with adjusted Rand index shown.

### 2.1 Set-based approach

*Scarecrow* creates two complementary sets of barcodes: (1) a ‘reference’ set for each whitelist of barcodes, and (2) a ‘mismatches’ set of all possible variants of each barcode on each whitelist allowing up to *m* nucleotide mismatches. Given an expected start position for a barcode of length *n*, an interval is defined that incorporates the jitter allowing barcoding searching within a range of ±*x* nucleotides up- and downstream. For example, for an expected start position of 10 with jitter = 2, the search range would span positions 8 to 12. For each position within this range, the *n*-length substring is extracted from the sequencing read and stored in a third ‘query’ set. In this example, the query set would contain five candidate sequences. The query set is compared to the whitelist reference set to first identify barcodes with exact matches. If any are found, these are returned in order of proximity to the expected start position. If no exact matches are found, the query set is then compared to the whitelist mismatches set. If any matches are found, these are returned in order of fewest mismatches first followed by closest distance to the expected start position. Accordingly, the *scarecrow* algorithm prioritises barcode matches in the following order: (1) a perfect match, (2) a match with the fewest nucleotide substitutions, (3) a match closest to the expected start position. If two or more barcodes satisfy all criteria equally (e.g. two candidates with one mismatch and both located one base from the expected start position), the result is treated as ambiguous, and a null match is returned. If jitter results in the search space range exceeding the start or end positions of a read, then the query sequence is trimmed at one end and padded with Ns at the other, enabling the above algorithm to proceed while treating the Ns as mismatches. If no valid matches are identified, *scarecrow* returns an invalid barcode – a string of N’s of length *n*.

### 2.2 Trie-based approach

The set-based matching approach is well-suited to relatively small barcode whitelists, such as 96 barcodes typically used at each indexing step in combinatorial barcoding protocols.

However, for larger barcode whitelists, such as those used with single-barcoding protocols, we have also implemented an alternative trie-based algorithm. During the *seed* step, this algorithm generates both a trie and a *k*-mer index from each whitelist of barcodes. The trie is used for exact barcode matching whilst the *k*-mer index supports approximate matching.

During the *reap* step, if no exact match is found during trie traversal, an approximate match is sought using the *k*-mer index. The same set of *n*-length ‘query’ sequences are extracted from the sequencing read, as described in the set-based approach. If the Hamming distance between the query sequence and a *k*-mer is ≤ maximum mismatches (user defined), then the *k*-mer is added to a list of candidate matches. Final barcode selection follows the prioritisation strategy as in the set-based method: exact match, fewest mismatches, and minimal distance from the expected start position.

The hybrid trie/*k*-mer approach is efficient because it significantly reduces the search space and does not require reference and mismatches sets to be held in memory. However, careful consideration must be given when setting the value of *k* as the *k*-mer needs a perfect match somewhere within the query sequence. For example, if the barcode length is 8 and *k*=4, then barcode ACGTTGCA will have the 4-mers ACGT, CGTT, GTTG, TTGC, TGCA. If we have a query sequence with two mismatches (denoted by lowercase letters), ACaTTaCA, then these mismatches impact all 4-mers and no match will be found. In such cases, a smaller value of *k*, e.g. *k* = 2, is required. We therefore define *k_max_* as the maximum *k* required to ensure at least one *k*-mer match: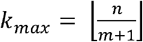, where *n* is the barcode length and *m* the allowed number of mismatches.

## 3 Workflow

We recommend that *seed* is run independently for each barcode whitelist, sampling 10,000 reads each time to generate well-supported barcode profiles. The outputs from *harvest* can be used to diagnose any potential issues with barcode incorporation or assumptions about library structure. For example, if a barcode returns a low read fraction (Figure S1) this can indicate that it was already demultiplexed by the sequencing provider. In such cases, the barcode should instead be recovered from the FASTQ header for each read using the *weed* command. To avoid redundant processing time, the corresponding barcode should be removed from the input barcode positions CSV before running *reap*. To ensure efficient barcode matching, barcode whitelists should contain sequences in the same orientation as they appear in the read. This is because it is computationally more efficient to match barcodes to reads in the same orientation than to reverse complement and re-search if no match is found. Finally, we recommend adapter trimming after running *scarecrow* to ensure consistent read lengths. This is because adaptor removal may lead to highly variable barcode positions, causing positional jitter to miss genuine barcodes outside of the expected range (e.g. ±2 nucleotides), resulting in loss of valid barcodes.

## 4 Examples

We illustrate the utility of *scarecrow* with publicly available data generated from Parse Biosciences Evercode and Scale Biosciences QuantumScale library preparations derived from (De Simone *et al.*, 2025)(Supplementary Material S2). These data are chosen because they profiled peripheral blood mononuclear cells, a diverse combination of immune cells, from the same donor across different commercial and widely used single-cell RNA-sequencing protocols. Considering the QuantumScale data, we show that applying jitter of 1 returns 65% more reads with matched (valid) barcodes compared to applying no jitter (n = 311,847,144 vs. 189,333,595; Figure 1B; Table S1). This also results in 67% more reads with perfectly matched barcodes (293,542,244 vs. 175,732,492), 65% more reads for alignment after sifting to remove reads with unmatched barcodes (311,768,984 vs.189,286,390), and 64% more uniquely mapped reads (214,478,807 vs. 130,506,898). These data gains propagate to downstream processing steps. After quantifying single-cell gene expression with *kallisto* and applying basic quality filtering, using jitter of 1 returned 50% more cells (12,789 compared to 8,530) and 5% more genes detected across cells (27,015 vs. 25,616) (Table S2). We also increased the median number of genes per cell with non-zero expression by 9% (744 compared to 684), a 10% increase in the median number of UMIs per cell (981 compared to 891), and a 21% increase in the maximum cell UMI observed (272 compared to 224) (Table S3). Therefore, recovering more reads at barcode error correction steps leads to higher-quality single-cell expression profiles for down-stream analysis tasks.

Our analyses indicate that approximately 2% of reads are assigned different barcodes when accounting for jitter (Table S4). This occurs because reads initially assigned to a barcode with a single mismatch with no jitter can be reassigned to perfect matching barcode with jitter > 0. To ensure that these reassignments did not introduce artifacts due to our advanced error correction, and to confirm the expected gains in data quality, we performed a uniform scRNA-seq analysis using *kallisto* (Melsted *et al.*, 2021) to generate count matrices with and without jitter applied on both Evercode and QuantumScale data sets. Following basic quality control to remove low quality cells and potential doublets, we identified common barcodes between the matrices generated with and without jitter. We then performed principal components analysis for dimensionality reduction, visualised the results using uniform manifold approximation and projection, and grouped cells into clusters using separate shared nearest neighbour graphs (Figure 1C). Comparing cluster membership highlighted the high concordance between both data sets (adjusted Rand index; ARI ≥ 0.94) indicating that our jitter-based barcode error correction did not negatively impact neighbour-finding or cell clustering. Moreover, we found no evidence of hybrid cells caused by barcode reassignment. In summary, *scarecrow* is a library agnostic pre-processing command-line tool that yields more reads and cells from combinatorial barcoding scRNA-seq data, enabling the maximal use of these highly scalable technologies.

