## Supplementary Material for "Harvesting more reads from single-cell combinatorial barcoding data with *scarecrow*"

### S1 Key functions of the *scarecrow* workflow

#### The *scarecrow* toolkit is comprised of multiple commands that execute the full workflow from barcode position identification through to assignment and barcode error correction. Additional functionality enables diverse single-cell combinatorial indexing libraries as input, including those using high throughput sequencing indices that are demultiplexed at the point of base calling, e.g. sci-RNA-seq3 (Cao *et al.*, 2019). The following section describes each command of the *scarecrow* workflow in detail.

#### S1.1 *seed*

This command accepts as its inputs a single set of FASTQ files in addition to a barcode whitelist file passed as a string with fields separated by colons (<barcode index>:<whitelist name>:<path to whitelist file>). The barcode whitelist is a text file with one barcode sequence per line and no header. Key optional parameters include: (1) the number of reads to sample, (2) an upper read count to sample from, (3) nucleotide frequency across reads that evidences a putative conserved sequence (linker base frequency), and (4) minimum length of conserved (linker) sequence. *seed* identifies highly frequent barcode positions and putative linker sequences within reads, returning a CSV file of barcode alignments, a TSV file of nucleotide frequencies for each FASTQ file, and a TSV file of conserved sequence runs. This step should be repeated for each barcode whitelist, for example the following bash code:

```
BARCODES=(BC1:n99_v5:${PROJECT}/barcode_whitelists/bc_data_n99_v5.txt
          BC2:v1:${PROJECT}/barcode_whitelists/bc_data_v1.txt
          BC3:v1:${PROJECT}/barcode_whitelists/bc_data_v1.txt)
FASTQS=(${PROJECT}/fastq/*.fastq.gz)
for BARCODE in ${BARCODES[@]}
do
    scarecrow seed \
        --num_reads 10000 \
        --upper_read_count 100000 \
        --fastqs ${FASTQS[@]} \
        --barcodes ${BARCODE} \
        --out ${PROJECT}/barcode_profiles/barcodes.${BARCODE%%:*}.csv
done
```

#### S1.2 *harvest*

The CSV files returned from *seed* for each barcode are passed to *harvest* to recover the barcode profiles. A TSV file of conserved sequences from *seed*, which will be identical across barcode whitelists, e.g. constant linker sequences, can also be provided to mask any barcode sequences that overlap these regions by chance. A minimum distance between barcodes can also be specified, in addition to the number of peaks (--barcode_count) to return per barcode whitelist. For example:

```
BARCODE_FILES=(${PROJECT}/barcode_profiles/barcodes.*.csv)
scarecrow harvest \
    ${BARCODE_FILES[@]} \
    --barcode_count 1 \
    --min_distance 10 \
    --conserved ${PROJECT}/barcode_profiles/barcodes.${BARCODES[0]%%:*}_conserved.tsv \
    --out ${PROJECT}/barcode_profiles/barcode_positions.csv
```

The predicted barcode positions are written to a CSV file, including the associated FASTQ file, barcode whitelist, orientation, and evidence in terms of read count and fraction:

barcode_whitelist,file_index,file,orientation,start,end,read_count,read_fraction

BC2:v1,2,SRR28867558_3.fastq.gz,forward,11,18,8852,0.93

BC3:v1,2,SRR28867558_3.fastq.gz,forward,49,56,8334,0.87

BC1:n99_v5,2,SRR28867558_3.fastq.gz,forward,79,86,7447,0.8

Histograms of the peaks identified from each FASTQ file and strand orientation are plotted to PNG files for inspection. The histograms can be informative for identifying potential issues with barcode matching, for instance if the sequencing reads have been demultiplexed by the sequencing provider leading to absent barcode peaks (Figure S1).

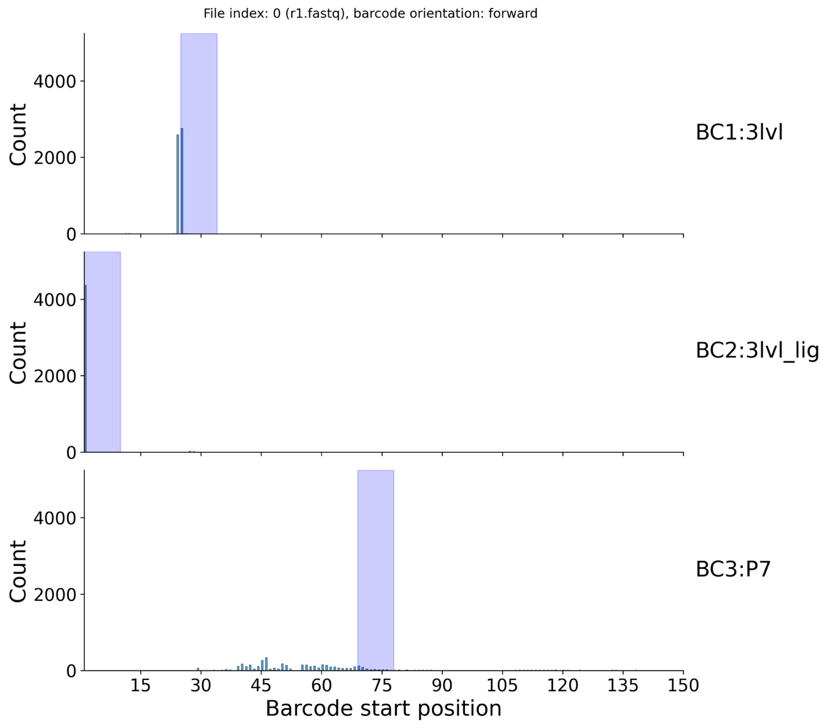

Figure S1. Histograms returned by *harvest* indicating good barcode matches for BC1 and BC2, and a poor match for BC3. This is a consequence of demultiplexing by the sequence provider, requiring BC3 to be extracted from the FASTQ read headers using *weed*.

#### S1.3 *reap*

This tool takes the barcode positions CSV generated by *harvest*, or a manually curated equivalent, along with the set of FASTQ files and barcode whitelists in the same format used by *seed*, and a target sequence range to extract (i.e. cDNA). Key options include: (1) the number of bases of jitter to consider, (2) maximum number of mismatches allowed during barcode matching, (3) minimum base quality, (4) output format (FASTQ or SAM), and (5) the UMI sequence range. For example:

```
THREADS=16
JITTER=1
MISMATCH=2
FASTQS=(${PROJECT}/fastq/*.fastq.gz)
OUT=$(basename ${FASTQS[0]%.fastq*})
mkdir -p ${PROJECT}/extracted/J${JITTER}M${MISMATCH}
sbatch -p uoa-compute --ntasks 1 --cpus-per-task ${THREADS} \
    --mem 16G --time=12:00:00 -o reap.%j.out -e reap.%j.err \
    scarecrow reap \
        --threads ${THREADS} \
        --batch_size 20000 \
        --fastqs ${FASTQS[@]} \
        --barcode_positions ${PROJECT}/barcode_profiles/barcode_positions.csv \
        --barcodes ${BARCODES[@]} \
        --extract 2:1-74 --umi 3:1-10 \
        --jitter ${JITTER} \
        --mismatch ${MISMATCH} \
        --out ${PROJECT}/extracted/J${JITTER}M${MISMATCH}/${OUT} \
        --out_fastq
```

The SAM output includes read tags for barcodes before (CR) and after (CB) correction, barcode start positions (XP) and mismatch counts (XM) for compatibility with downstream tools. The FASTQ output an interleaved FASTQ file, the first read of which contains the barcodes and UMI sequence, the second read contains the target sequence. The read headers in the FASTQ file incorporate the SAM read tag information. In addition, the FASTQ output also generates a JSON file that includes the parameters required to use the FASTQ files with the *kallisto-bustools* (Sullivan *et al.*, 2025) *kb count* tool. A file summarising the number of reads for each possible mismatch count across the barcodes is also generated, along with a file summarising the number of reads containing barcodes starting at each possible position accounting for jitter.

#### S1.4 Ancillary functions

- ***weed***. If the data have been demultiplexed by the sequencing provider, this command can append a specified sequence index from the FASTQ read header to the associated read CB tag in a SAM file generated by *reap*. Barcode correction is performed as per *reap* and so jitter and mismatch values can be set.
- ***sift***. The SAM or FASTQ file generated by reap can be filtered to remove reads that contain invalid barcodes.
- ***recast***. This command converts between the *scarecrow* SAM and FASTQ files, and *vice versa*.
- ***stats*** accepts a *scarecrow* SAM or FASTQ file and generates count data on the following read tags: (1) uncorrected barcode sequence at the expected position (CR), (2) corrected barcode sequence accounting for jitter (CB), (3) corrected barcode start position (XP), (4) corrected barcode mismatch count (XM), and (5) UMI sequences (UR). These count data can be summarised over input files for inspection, exploratory data analysis and quality control.

### S2 Examples

#### S2.1 Scale Biosciences QuantumScale

QuantumScale data was downloaded from the NCBI Sequence Read Archive (SRA accession: SRR28867557). Code is available on processing this data with *scarecrow* at:

<https://github.com/MorganResearchLab/scarecrow/blob/main/docs/example_scale.md>

#### S2.2 Parse Evercode (WTv2)

Evercode WTv2 data was downloaded from the SRA (accession: SRR28867558). Code is available on processing this data with *scarecrow* at:

<https://github.com/MorganResearchLab/scarecrow/blob/main/docs/example_evercode.md>

#### S2.3 Analysis

The code to analyse the data and generate the figures presented here is available at:

<https://github.com/MorganResearchLab/scarecrow_paper>

The number of reads with valid (matched) barcodes identified by *scarecrow* at different jitter values is shown in Table S1, along with the number of reads with perfect barcode matches, and the total barcode count. We observe a 12% increase in the number of reads with valid barcodes when applying jitter *j* = 2 to the Evercode v2 data, and a 65% increase when applying *j* = 1 to the QuantumScale data, relative to applying no jitter (*j* = 0). The number of perfect barcodes identified, i.e. those with zero mismatches, increased by 6% at *j* = 2 in the Evercode data, and 67% at *j* = 1 in the QuantumScale data. The total number of barcode combinations (BC1_BC2_BC3) identified increased by 2% at *j* = 2 in the Evercode data, and by 9% at *j* = 1 in the QuantumScale data. After sifting the reads output by *scarecrow* to remove invalid (unmatched) barcodes, and trimming to remove adapter sequences, there was a 12% increase in reads input to alignment at *j* = 2 in the Evercode data, and a 65% increase at *j* = 1 in the QuantumScale data. This culminated in an increase in the number of uniquely mapped reads of 11% in the Evercode data and 64% in the QuantumScale data.

Table S1. Summary of barcodes identified by *scarecrow* and uniquely mapped reads by *STAR*

Reads with valid and perfect barcodes matched by scarecrow, in addition to the number of barcodes identified are reported. The number of input reads for alignment is post-trimming adapter sequences and removal of reads with invalid barcodes. The number of uniquely mapped reads is reported by STAR.

|  |  | ***scarecrow*** | | | **STAR** | | |
| --- | --- | --- | --- | --- | --- | --- | --- |
| **Dataset** | **Jitter** | **Valid barcodes** | **Perfect barcodes** | **Barcode count** | | **Input reads** | **Uniquely mapped** |
| Evercode | 0 | 132,932,347 | 119,851,052 | 880,650 | | 132,869,473 | 90,975,102 |
|  | 1 | 145,661,189 | 125,784,335 | 894,238 | | 145,489,281 | 98,926,170 |
|  | 2 | 148,654,886 | 126,768,927 | 898,204 | | 148,409,133 | 100,771,140 |
| QuantumScale | 0 | 189,333,595 | 175,732,492 | 270,459 | | 189,286,390 | 130,506,898 |
|  | 1 | 311,847,144 | 293,542,244 | 293,898 | | 311,768,984 | 214,478,807 |

After processing each dataset with *scarecrow*, and generating *kallisto* count matrices at jitter = [0,1] for QuantumScale data and jitter = [0, 1, 2] for the Evercode data, the unfiltered count matrices were analysed. For a given dataset, the count matrices for each jitter *j* setting were processed to: (1) check for library saturation; (2) identify the UMI count inflection point (min UMIs = 100); (3) apply a basic cell (min UMIs = 100) and feature (min features = 1) count filters; and (4) plot distributions of features, cells, and mitochondrial (mt) gene percentage.

We show that there is little change in library saturation between different jitter settings (QuantumScale: *j* = 0, Figure S2A; *j* = 1, Figure S3A; Evercode: *j* = 0, Figure S4A; *j* = 2, Figure S5A). There is an increase in the number of cells with at least 100 UMIs when applying jitter, which is more notable in the QuantumScale data (*j = 0,* Figure S2B, n = 8530; *j* = 1, Figure S3B, n = 12789) than the Evercode data (*j = 0,* Figure S4B, n = 16379; *j* = 2, Figure S5B, n = 16420). There is little difference in the distributions of features, cells, and percentage mt genes with or without the application of jitter (QuantumScale: *j* = 0, Figure S2C; *j* = 1, Figure S3C; Evercode: *j* = 0, Figure S4C; *j* = 2, Figure S5C). The density of the percentage mitochondrial genes relative to UMI count also shows little difference with or without the application of jitter (QuantumScale: *j* = 0, Figure S2D; *j* = 1, Figure S3D; Evercode: *j* = 0, Figure S4D; *j* = 2, Figure S5D). These results indicate that applying jitter when undertaking barcode matching increases the usable data without adversely affecting the underlying distributions of cells, features, and mitochondrial gene percentage.

A summary of the barcode, UMI and feature counts for each *kallisto* count matrix is provided in Table S2. Comparing the counts with and without the application of jitter, indicates that pre-filtering, the QuantumScale data returned a 33% increase in barcode count at *j* = 1, 64% more UMIs, and 5% more total features detected. While post-filtering, 50% more cells were retained than at *j* = 0. The Evercode data returned a 7% increase in barcode count at *j* = 2, 11% more UMIs, and a 1% increase in total features. Post-filtering resulted in 0.25% more cells than at *j* = 0. The average number of genes and UMIs per cell post-filtering is reported in Table S3, along with the maximum UMIs observed across cells. These values translate to a 9% increase in genes per cell and 10% increase in UMIs per cell in the QuantumScale data at *j* = 1 compared to *j* = 0, and a 6% increase in genes per cell and 8% increase in UMIs per cell in the Evercode data at *j* = 2 compared to *j* = 0. We also observe between a 6-21% increase in the maximum gene UMI count observed across cell.

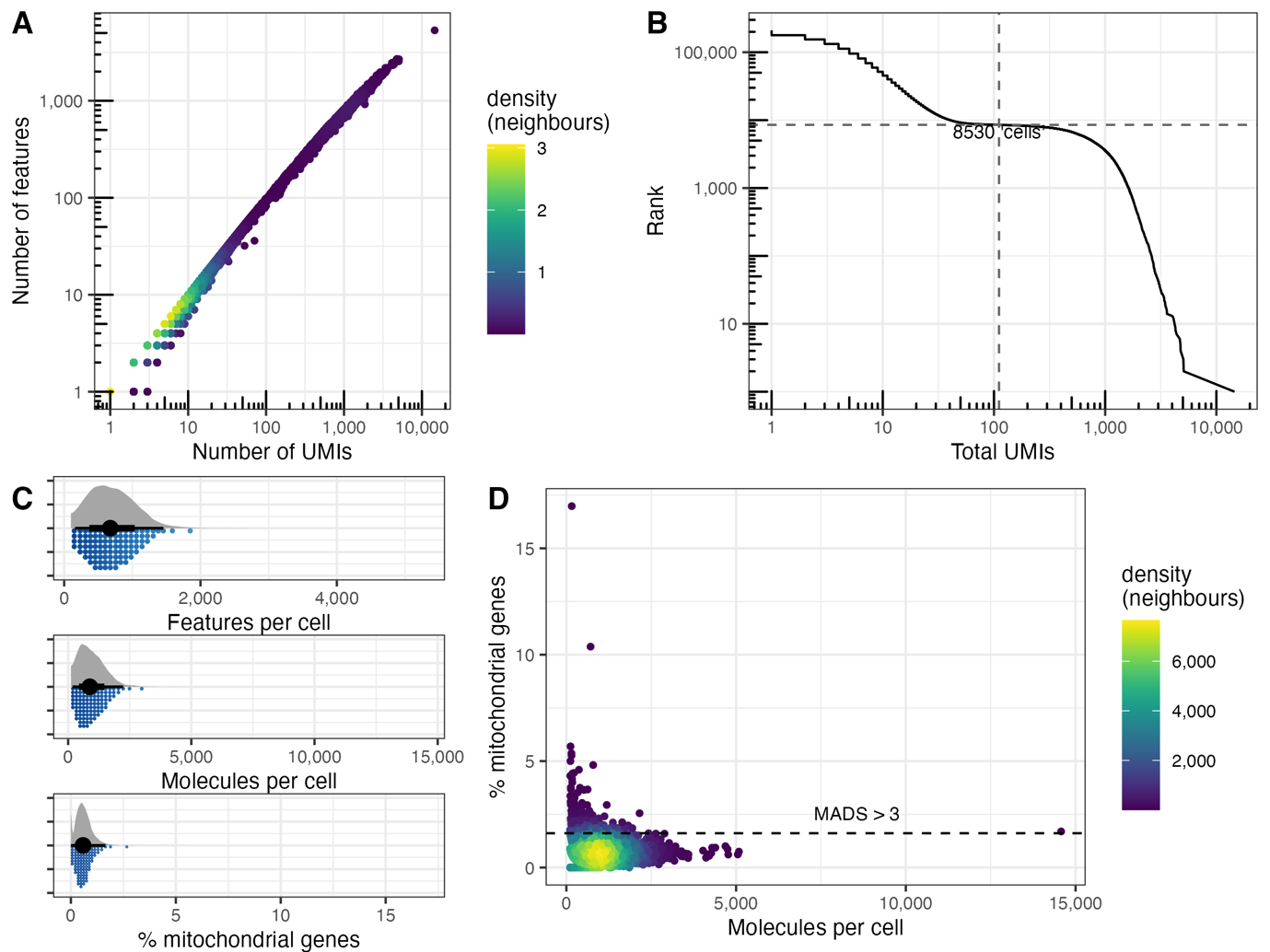

Figure S2. Summary metrics of QuantumScale data processed with scarecrow at jitter 0 followed by count matrix generation with kallisto

(A) Library saturation density plot showing number of molecules against number of features. (B) Knee plot showing inflection point for number of cells with at least 100 UMIs. (C) Rain cloud distribution plots of features per cell, molecules per cell, and % mitochondrial genes. (D) Density plot showing molecules per cell against % mitochondrial genes, with horizontal line indicating 99^th^ percentile of % mitochondrial genes.

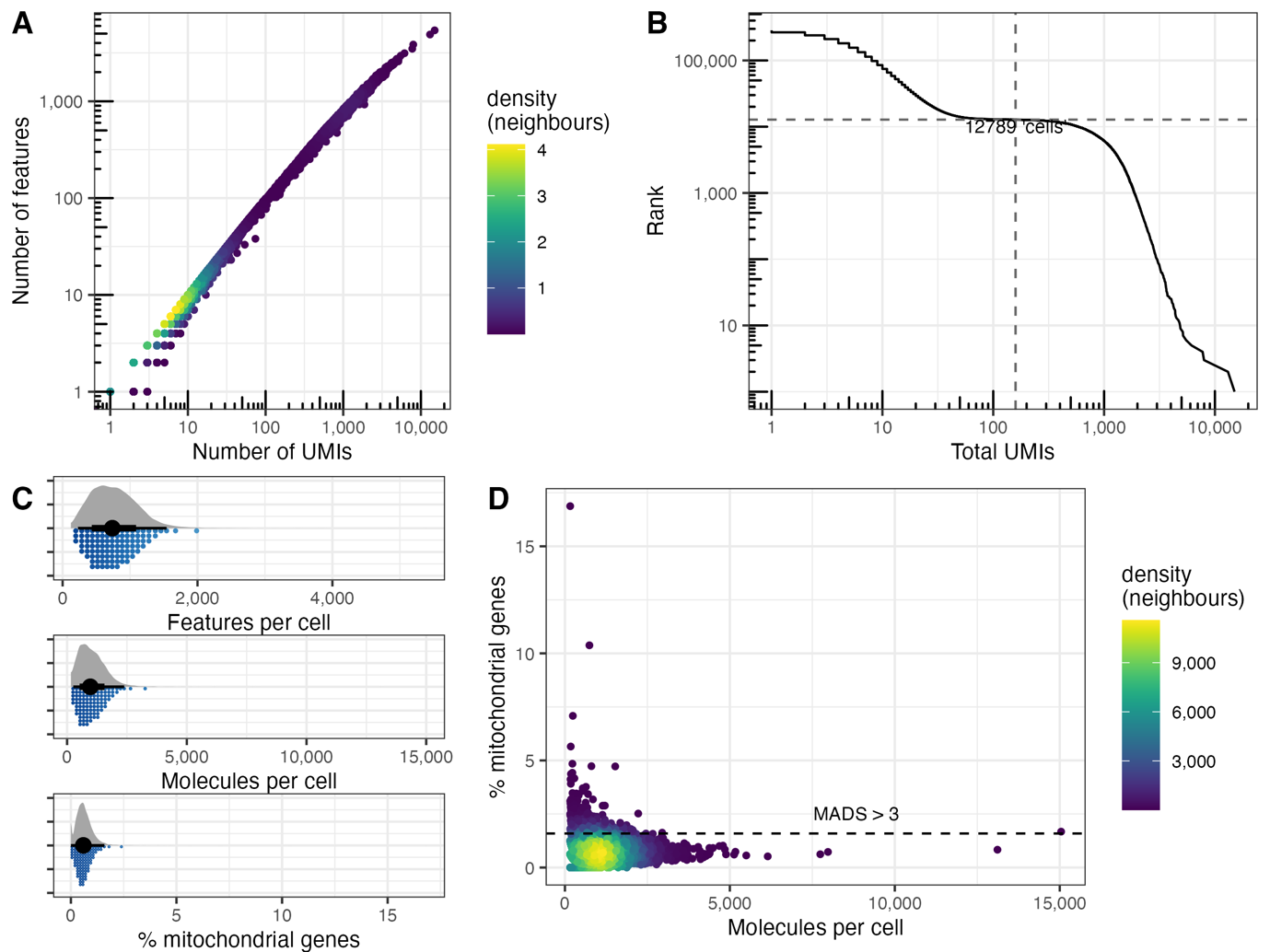

Figure S3. Summary metrics of QuantumScale data processed with *scarecrow* at jitter 1 followed by count matrix generation with *kallisto*

(A) Library saturation density plot showing number of molecules against number of features. (B) Knee plot showing inflection point for number of cells with at least 100 UMIs. (C) Rain cloud distribution plots of features per cell, molecules per cell, and % mitochondrial genes. (D) Density plot showing molecules per cell against % mitochondrial genes, with horizontal line indicating 99^th^ percentile of % mitochondrial genes.

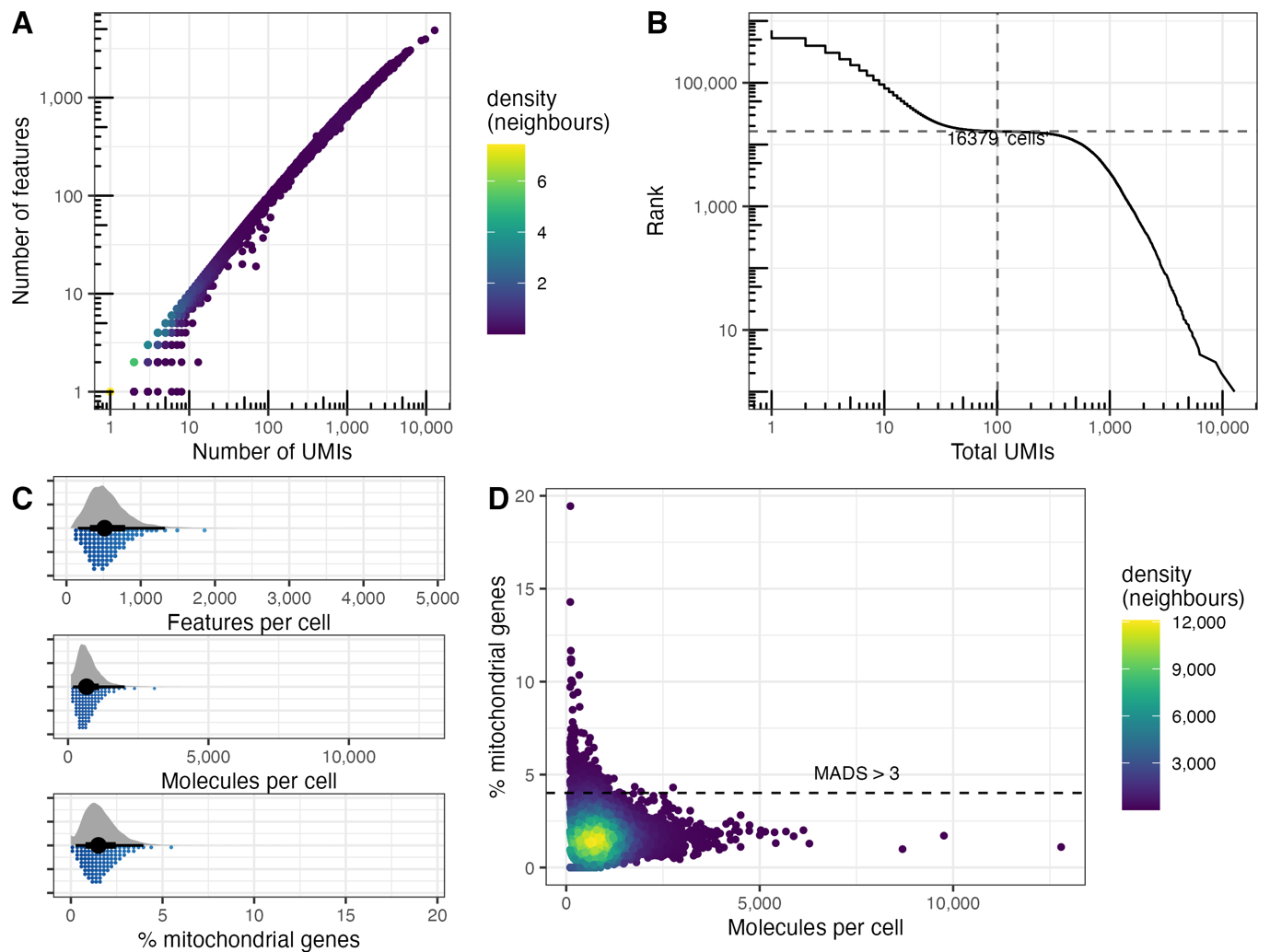

Figure S4. Summary metrics of Evercode data processed with *scarecrow* at jitter 0 followed by count matrix generation with *kallisto*

(A) Library saturation density plot showing number of molecules against number of features. (B) Knee plot showing inflection point for number of cells with at least 100 UMIs. (C) Rain cloud distribution plots of features per cell, molecules per cell, and % mitochondrial genes. (D) Density plot showing molecules per cell against % mitochondrial genes, with horizontal line indicating 99^th^ percentile of % mitochondrial genes.

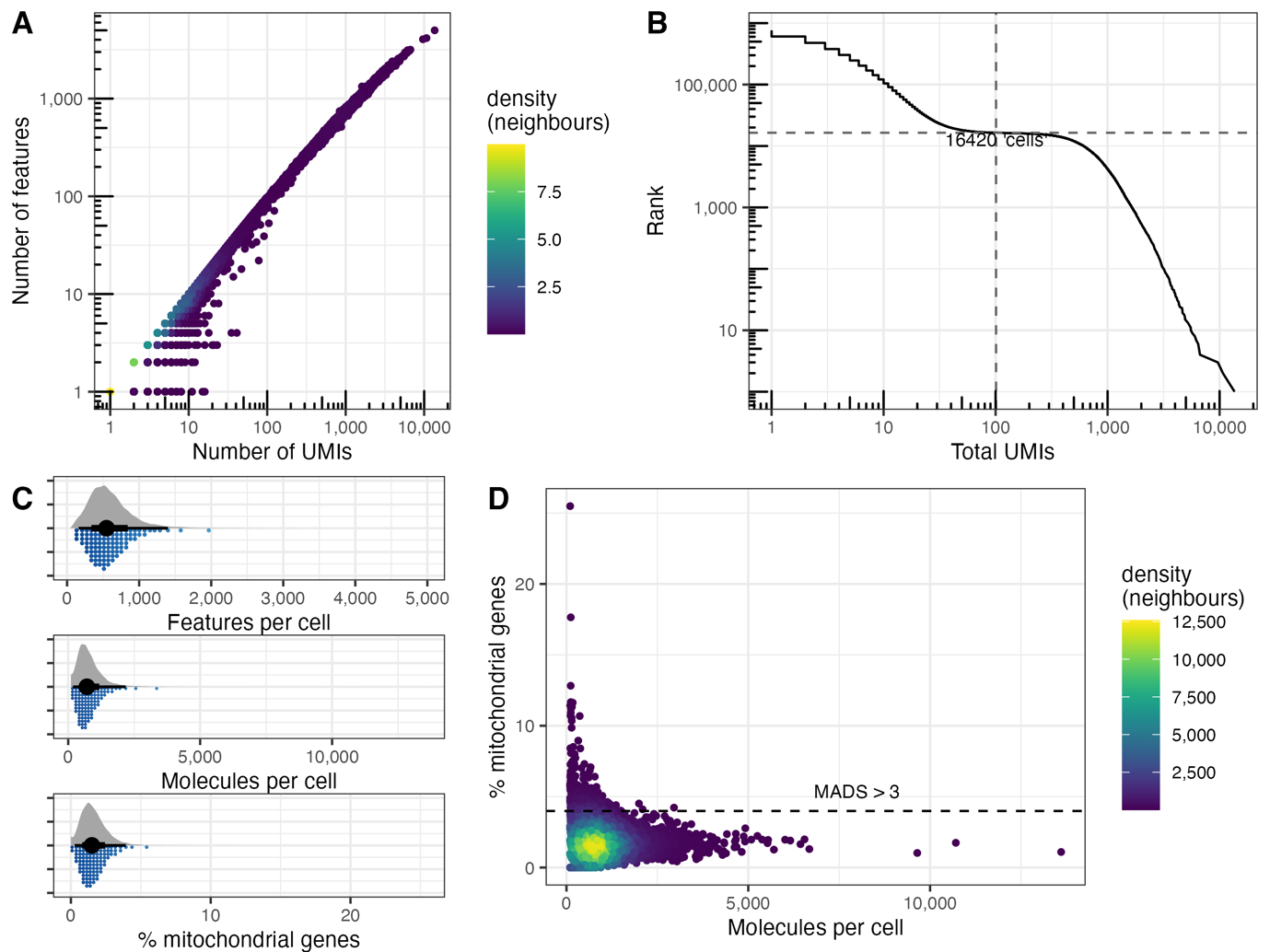

Figure S5. Summary metrics of Evercode data processed with *scarecrow* at jitter 2 followed by count matrix generation with *kallisto*

(A) Library saturation density plot showing number of molecules against number of features. (B) Knee plot showing inflection point for number of cells with at least 100 UMIs. (C) Rain cloud distribution plots of features per cell, molecules per cell, and % mitochondrial genes. (D) Density plot showing molecules per cell against % mitochondrial genes, with horizontal line indicating 99^th^ percentile of % mitochondrial genes.

Table S2. Summary of counts

The *kallisto* barcode, UMI, and feature counts are shown pre-filtering. The number of cells remaining after filtering on the inflection point and mitochondrial gene percentage are also shown, the number of features remains unchanged post-filtering based on the inflection point and subsequent mitochondrial gene percentage.

|  |  | **kallisto** | | **Features post-filter** | **Cells post-filter** | |
| --- | --- | --- | --- | --- | --- | --- |
| **Dataset** | **Jitter** | **Barcode count** | **UMI count** |  | **Cells** | **mt < 0.99** |
| Evercode | 0 | 755,765 | 15,734,784 | 27,207 | 16,379 | 16,002 |
|  | 1 | 793,222 | 17,134,428 | 27,372 | 16,397 | 16,017 |
|  | 2 | 806,170 | 17,469,472 | 27,395 | 16,420 | 16,037 |
| QuantumScale | 0 | 215,567 | 9,624,059 | 25,616 | 8,530 | 8,298 |
|  | 1 | 287,665 | 15,746,814 | 27,015 | 12,789 | 12,477 |

Table S3. Average number of genes and UMIs per cell in post-filtered data.

| **Dataset** | **Jitter** | **Median genes/cell** | **Median UMIs/cell** | **Max UMIs/gene** |
| --- | --- | --- | --- | --- |
| Evercode | 0 | 519 | 664 | 273 |
|  | 1 | 549 | 712 | 289 |
|  | 2 | 553 | 716 | 289 |
| QuantumScale | 0 | 684 | 891 | 224 |
|  | 1 | 744 | 981 | 272 |

Plotting the mean and max counts per gene at jitter = 0 against jitter = 1 in the QuantumScale data (Figure S6), and at jitter = 0 against jitter = 2 in the Evercode data (Figure S7), indicates that applying jitter typically results in an increase in both metrics. However, this is not always the case, suggesting that some reads are assigned to different cells. This is possible because *scarecrow* can both identify a closer matching barcode when using jitter > 0, and identify multiple best-matches taking into account both jitter and sequencing errors; the latter results in a null/invalid barcode being returned.

To investigate read-barcode re-assignment in depth, we first compared the counts with and without jitter for each unique combination of barcodes reported by *scarecrow stats* (Figures S8 and S9). Most barcodes show a higher UMI count when jitter is applied (positive increase in ΔUMI; QuantumScale slope = 1.619; Evercode slope = 0.041). The negative ΔUMI for some barcodes in the intersect indicates the assignment of reads to different barcodes. To further explore this, we sampled 10,000 unique barcodes from the jitter = 0 results and identified the sequencing reads associated with these barcodes. The barcode assignments for these same reads were then retrieved from the results with jitter applied (QuantumScale jitter = 1; Evercode jitter = 2), and the difference in barcode UMI counts compared (Figures S8 and S9).

We observed a 1.8-2.5% difference in barcode assignment when accounting for jitter (Table S4). The reads associated with the 10,000 sampled barcodes returned 11,199 barcodes at jitter = 1 in the QuantumScale data, and 15,693 barcodes at jitter = 2 in the Evercode data. A small (<1%) subset of reads did not return a valid barcode when jitter was applied (QuantumScale n = 242; Evercode *n* = 914). On further investigation, this was due to ambiguity in barcode assignment. For example, when accounting for jitter, on rare occasions more than one barcode can be perfectly matched and equidistant from the expected barcode position. In such instances it is not possible to determine which match is most likely, and the barcode is recorded as missing. We found that 92% of barcodes re-assigned at jitter = 1 in the QuantumScale data were perfectly matched to a barcode on a whitelist, whereas perfect matches were found for only 54% of barcodes re-assigned at jitter = 2 in the Evercode data. The mean number of mismatches for each barcode when accounting for jitter was equal to or less than applying no jitter. Notably, most mismatches were found on the barcode in the reverse transcription step (BC3), indicating a limitation in combinatorial indexing protocols that relies on reverse transcriptase to incorporate DNA barcodes.

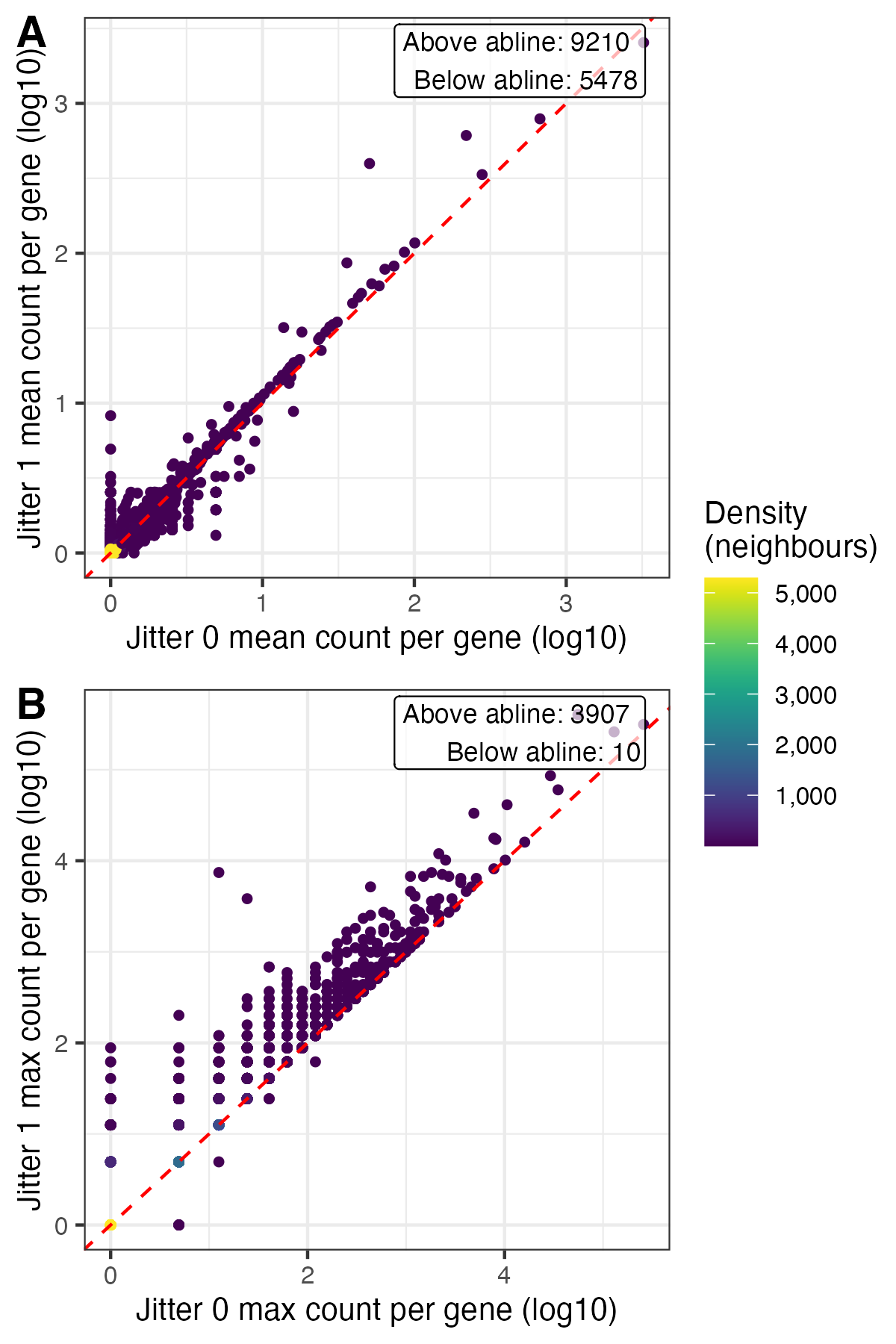

Figure S6. Density scatter plots based on filtered *kallisto* count matrices of QuantumScale data

(A) Shows mean UMI count per gene. Points above the line indicate genes with an increase in UMI count at jitter = 1 relative to jitter = 0, while those below the line indicate a decrease in UMI count. (B) Shows an equivalent analysis for max UMI count per gene.

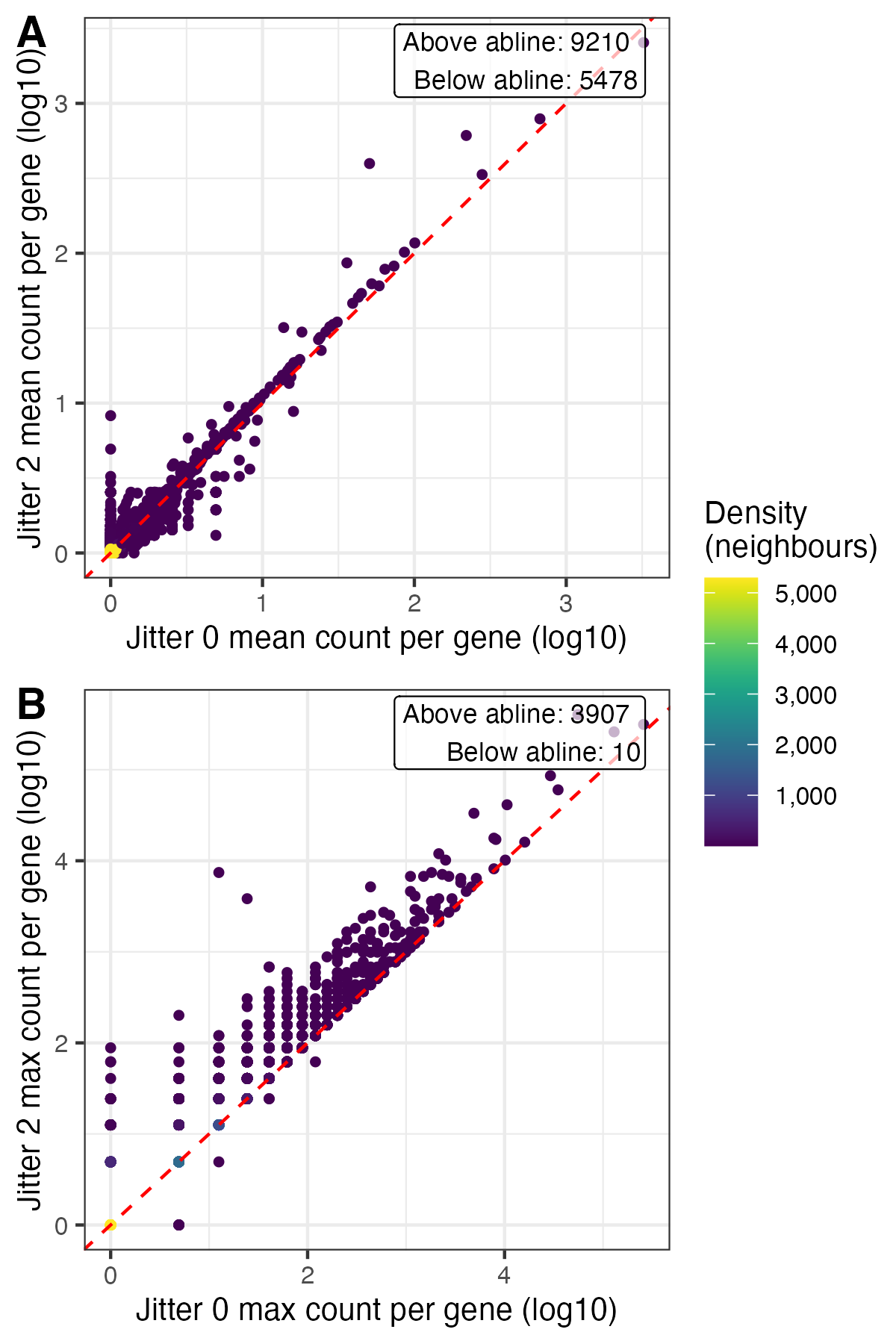

Figure S7. Density scatter plots based on filtered *kallisto* count matrices of Evercode data

(A) Shows mean UMI count per gene. Points above the line indicate genes with an increase in UMI count at jitter = 2 relative to jitter = 0, while those below the line indicate a decrease in UMI count. (B) Shows an equivalent analysis for max UMI count per gene.

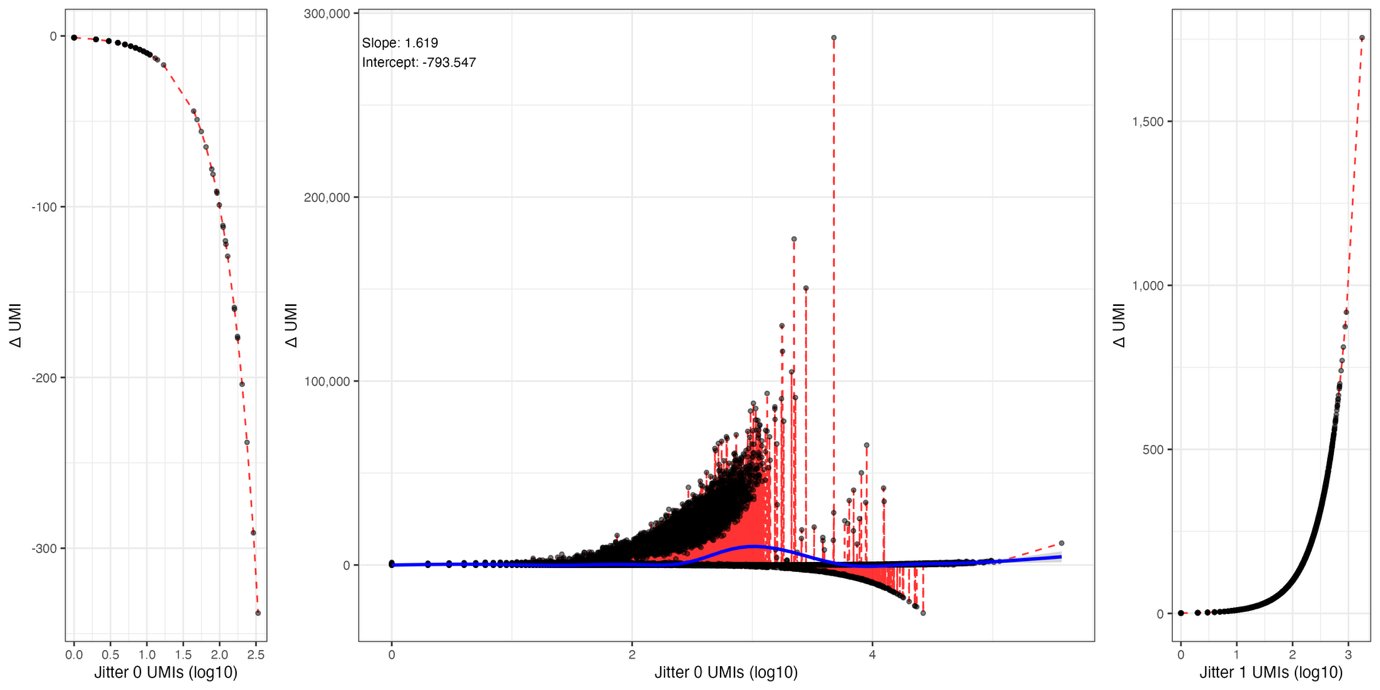

Figure S8. Barcode UMI count difference between jitter 0 and jitter 1 in the QuantumScale data. Difference (Δ) is calculated in each panel as the barcode UMI count at jitter 1 – the barcode UMI count at jitter 0. The left panel shows barcode UMI counts unique to jitter = 0 plotted against the difference in counts relative to jitter 1. As these barcodes are unique to jitter 0, Δ is negative. The right panel shows the equivalent for barcode UMI counts unique to jitter = 1, hence the Δ is positive. The middle plot shows Δ for the intersect, barcodes identified at jitter = 0 and jitter = 1.

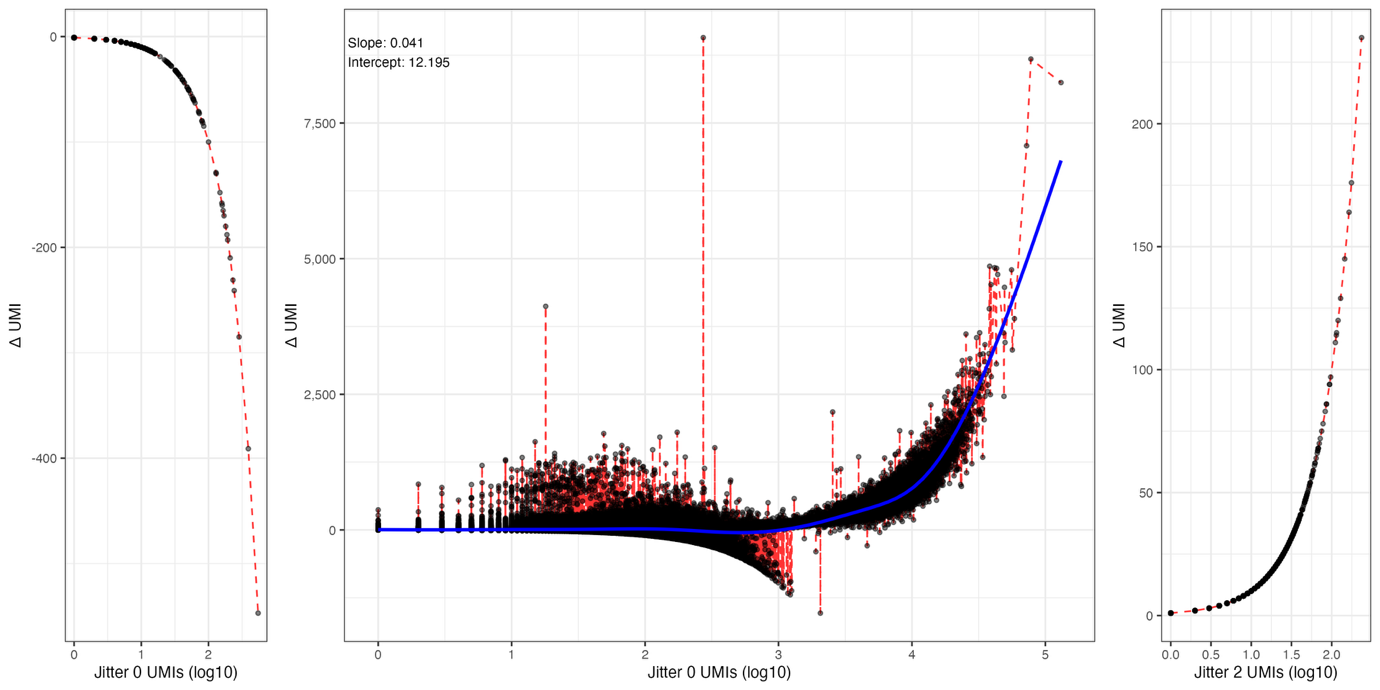

Figure S9. Barcode UMI count difference between jitter 0 and jitter 2 in the Evercode data. Difference (Δ) is calculated in each panel as the barcode UMI count at jitter 2 – the barcode UMI count at jitter 0. The left panel shows barcode UMI counts unique to jitter = 0 plotted against the difference in counts relative to jitter 2. As these barcodes are unique to jitter 0, Δ is negative. The right panel shows the equivalent for barcode UMI counts unique to jitter = 2, hence the Δ is positive. The middle plot shows Δ for the intersect, barcodes identified at jitter = 0 and jitter = 2.

Table S4. Analysis of sequencing read barcode assignment in 10,000 sampled barcodes

For each dataset, 10,000 unique barcodes were sampled from the results without jitter. The reads associated with these barcodes were then identified from the results where jitter was applied, and their barcode assignments compared to those without jitter. The table indicates: the number of unique barcodes identified across reads with and without the application of jitter; the number of reads with barcodes that were identical, different, or missing (N/A) when accounting for jitter; among the subset of reads with re-assigned barcodes, the number of reads with and without mismatches in their assigned barcodes; the mean number of mismatches across reads for each barcode.

|  |  | **QuantumScale** | | **Evercode** | |
| --- | --- | --- | --- | --- | --- |
|  | Jitter | 0 | 1 | 0 | 2 |
|  | Barcodes | 10,000 | 11,199 | 10,000 | 15,693 |
| Barcode assignment | Identical |  | 7,482,707 (98.1%) |  | 1,220,122 (97.5%) |
|  | Different |  | 140,884 (1.8%) |  | 30,957 (2.5%) |
|  | N/A |  | 242 (<1%) |  | 914 (<1%) |
| Barcode re-assignment mismatches | 0 |  | 129,833 (92.2%) |  | 16,619 (53.7%) |
|  | >0 |  | 11,051 (7.8%) |  | 14,338 (46.3%) |
| Mean mismatches | BC1 | 0.02 | 0.02 | 0.03 | 0.03 |
|  | BC2 | 0.02 | 0.02 | 0.05 | 0.03 |
|  | BC3 | 0.09 | 0.05 | 0.11 | 0.08 |

To explore if the application of jitter leads to incongruous barcode assignments leading to hybrid cells, we conducted a series of *k*-nearest neighbour analyses comparing QuantumScale jitter = 0 to jitter = 1, and Evercode jitter = 0 to jitter = 2. Raw count matrices were filtered to retain cells and genes with at least 100 counts each. Within each dataset (QuantumScale, Evercode), the filtered matrices were subset to ensure matching genes and barcodes between the jitter values. The *k*-nearest neighbours were then identified for *k* values of 2:10, 20, 30, and 50 using the fast nearest neighbour method (R FNN package). For each *k* value, we calculated the proportion of shared neighbours for each cell (Figures S10 and S11). The mean proportion of shared neighbours increased with increasing *k* values, ranging from 77-88% in the QuantumScale data and 61-77% in the Evercode data (Table S5).

We further explored any possible impact on the application of jitter on barcode matching by processing the data in a typical workflow. The raw count matrices generated with and without the application of jitter were filtered to remove genes with zero counts and retain cells with at least 500 UMIs and fewer than three median absolute deviations in mitochondrial gene content. Doublets were identifed using the R package *scDblFinder* (Germain *et al.*, 2022) taking the union of doublets labelled by either the densities or classification approaches. Singlets were extracted from the filtered matrices for genes and barcodes common to both. Using the same random seed on each matrix, gene expression counts were log-normalised, a PCA run, a shared nearest-neighbour graph constructed, cluster assignments made based on the Louvain algorithm, and a UMAP plot generated. The clustering results of the QuantumScale Bio data at jitter = 1 compared to jitter = 0 (Figure S12), and the Evercode data at jitter = 2 compared to jitter = 0 (Figure S13), returned adjusted Rand indices of 0.98 and 0.94, respectively, indicating near-perfect agreement. A less stringent QC, repeating the analysis on cells with at least 100 UMIs, returned adjusted Rand indices of 0.93 and 0.97, respectively.

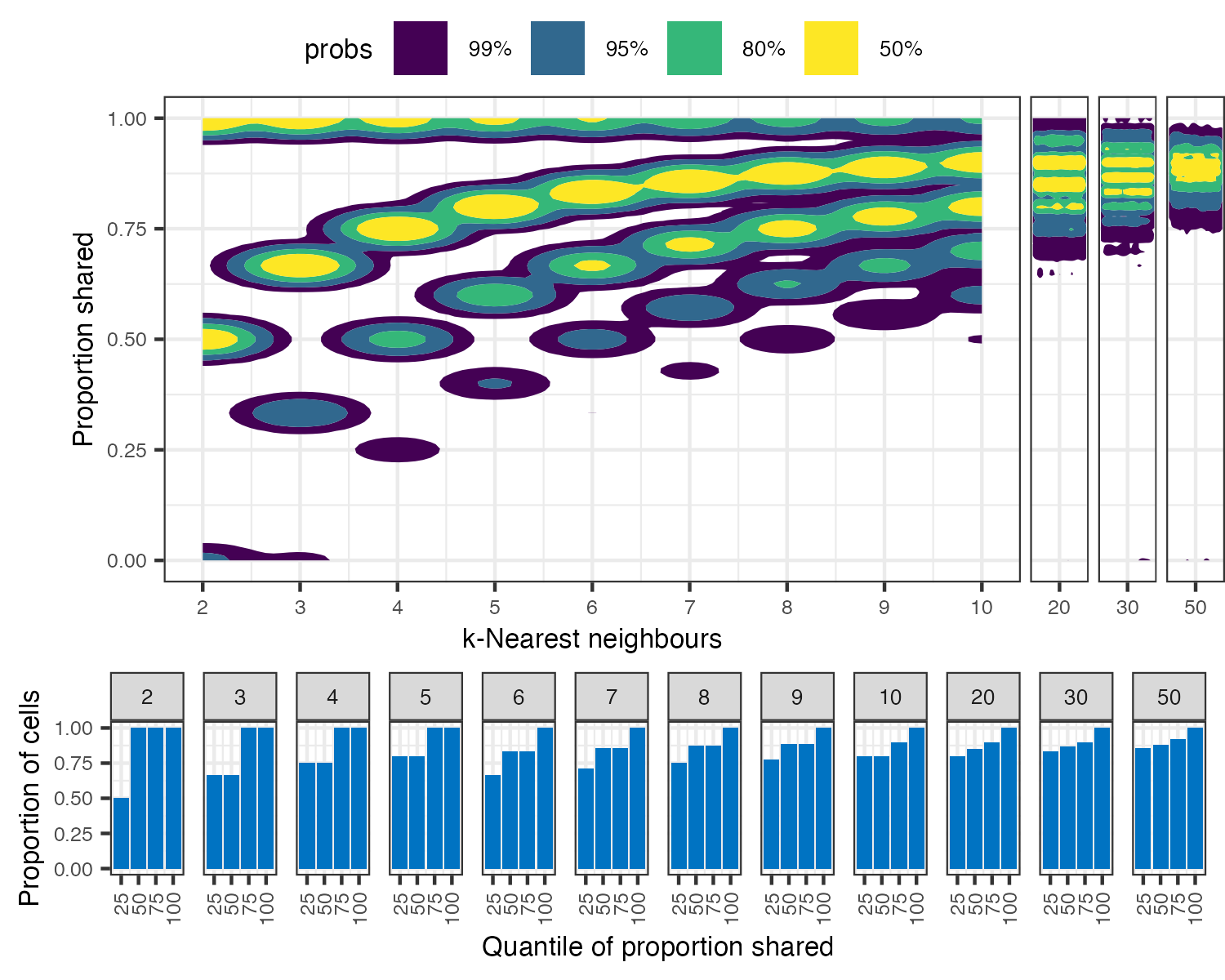

Figure S10. *k*-Nearest neighours analysis of QuantumScale data for jitter = 0 against jitter = 1. Plot of high-density regions estimates of proportion of shared neighbours at different *k* values. A small amount of jitter (1 x 10^-8^) has been applied to x axis (*k*) values to facilitate 2D density estimations. Bar plots illustrate the proportion of cells for each quantile of shared neighbours.

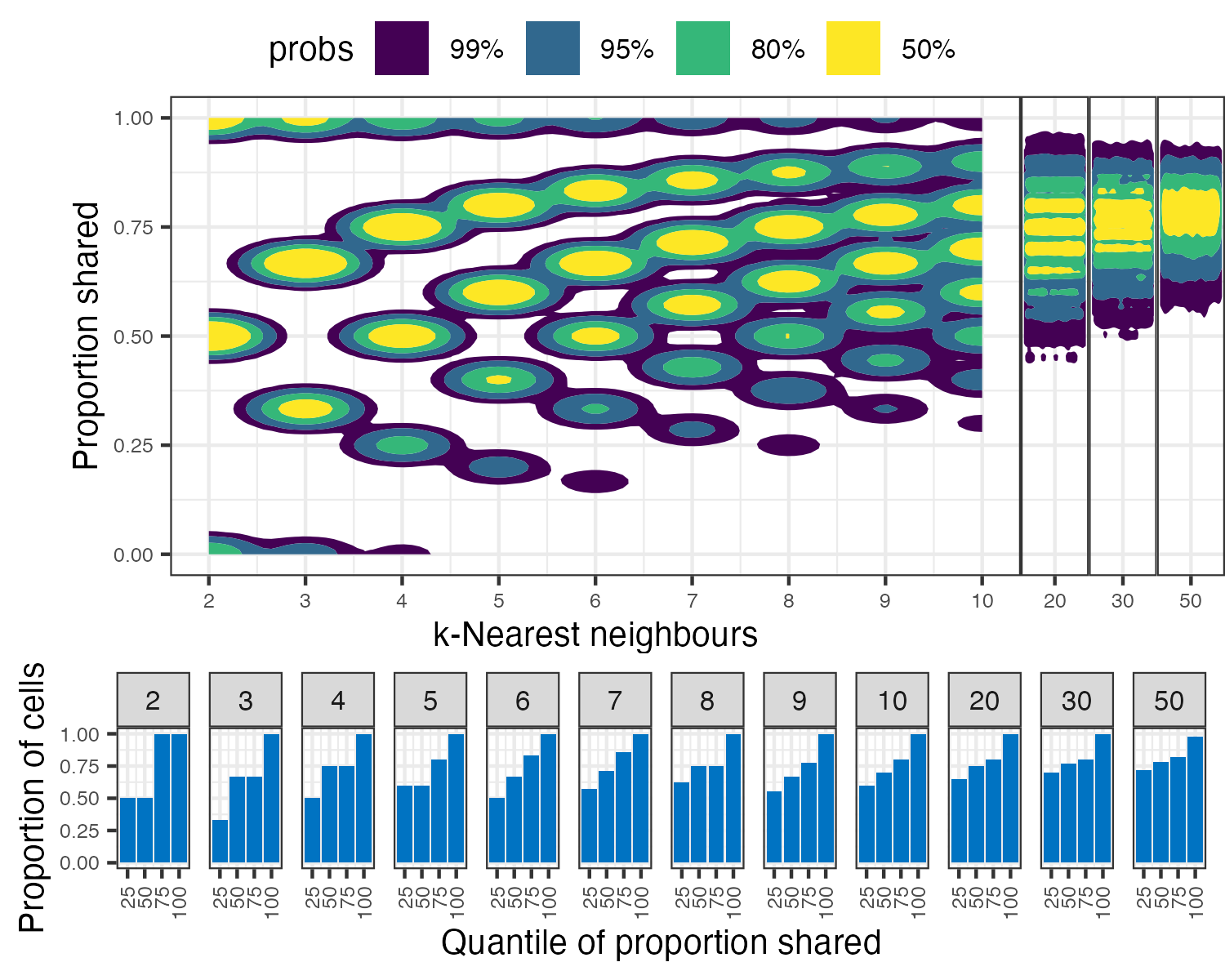

Figure S11. *k*-Nearest neighours analysis of Evercode data for jitter = 0 against jitter = 2. Plot of high-density regions estimates of proportion of shared neighbours at different *k* values. A small amount of jitter (1 x 10^-8^) has been applied to x axis (*k*) values to facilitate 2D density estimations. Bar plots illustrate the proportion of cells for each quantile of shared neighbours.

Table S5. Mean proportion of shared neighbours at varying *k* values

| ***k*** | **QuantumScale** | **Evercode** |
| --- | --- | --- |
| 2 | 0.772 | 0.609 |
| 3 | 0.785 | 0.635 |
| 4 | 0.798 | 0.650 |
| 5 | 0.805 | 0.662 |
| 6 | 0.814 | 0.673 |
| 7 | 0.819 | 0.680 |
| 8 | 0.823 | 0.687 |
| 9 | 0.83 | 0.693 |
| 10 | 0.834 | 0.699 |
| 20 | 0.856 | 0.731 |
| 30 | 0.866 | 0.749 |
| 50 | 0.879 | 0.772 |

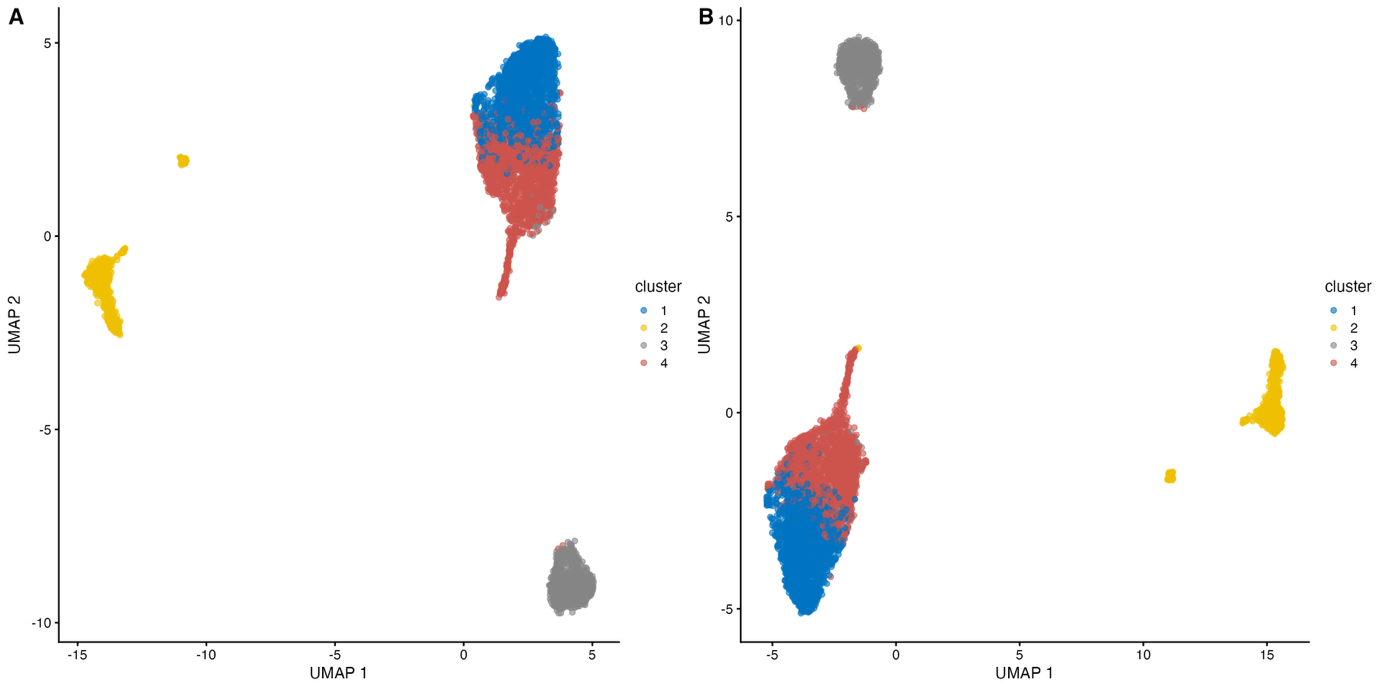

Figure S12. UMAP of QuantumScale data for jitter = 0 (A) against jitter = 1 (B)

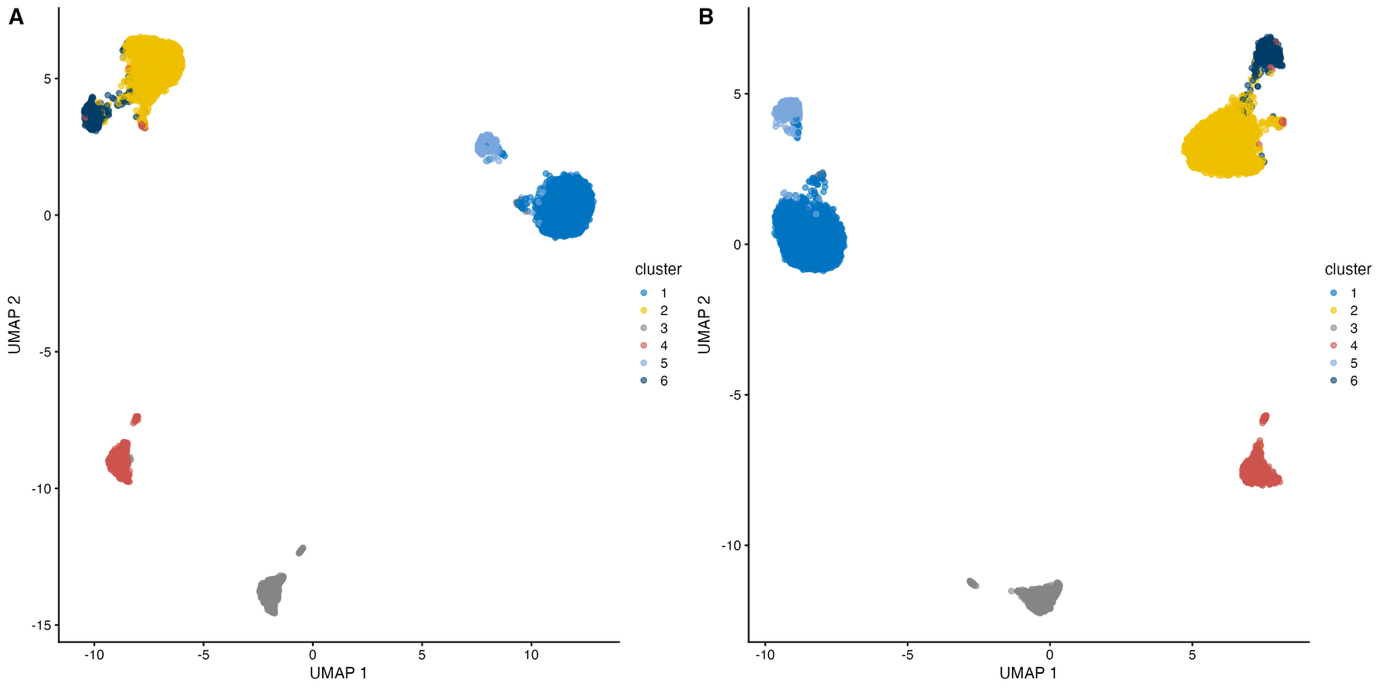

Figure S13. UMAP of Evercode data for jitter = 0 (A) against jitter = 2 (B)
